# A Wide-Field Micro-Computed Tomography Detector: Micron Resolution at Half-centimeter Scale

**DOI:** 10.1101/2021.08.27.457808

**Authors:** Maksim A. Yakovlev, Daniel J. Vanselow, Mee Siing Ngu, Carolyn R. Zaino, Spencer R. Katz, Yifu Ding, Dula Parkinson, Steve Yuxin Wang, Khai Chung Ang, Patrick J. La Riviere, Keith C. Cheng

## Abstract

Ideal 3-dimensional imaging of many complex samples, such as biological tissues made up of micro-scale structures extending over millimeter- to centimeter-scale tissue samples and organisms, requires both a wide field-of-view and high resolution. With existing optics and detectors used for micro-CT imaging, sub-micron pixel resolution can only be achieved for fields-of-view of <2 mm. This manuscript presents a unique detector system with a 6-mm field-of-view image circle and 0.5 μm pixel size that can be used in both synchrotron facilities and tabletop micro-CT units. A resolution-test pattern with linear microstructures and whole adult *Daphnia magna* were imaged on Beamline 8.3.2 of the Advanced Light Source. Volumes of 10,000 × 10,000 × 7,096 isotropic 0.5 μm voxels were reconstructed over a 5.0 × 3.5 mm field-of-view. Measurements in the projection domain confirmed a 1.182 μm measured spatial resolution that is largely Nyquist-limited. This unprecedented combination of field-of-view and resolution dramatically reduces the need for sectional scans and computational stitching for large samples, ultimately offering the means to elucidate change in tissue and cellular morphology in the context of larger whole, intact model organisms and specimens. This development is also anticipated to benefit micro-CT imaging in materials science, microelectronics, agricultural science, and biomedical engineering.

**Synopsis:** A custom wide-field lens and a new-generation megapixel camera enabled microCT scanning over a 3.5 × 5 mm field-of-view at a 1 μm resolution / 0.5 μm pixel size at the Berkeley Lawrence Advanced Light Source using a phantom with micron scale features. This novel combination of resolution and field-of-view will be broadly applicable to any setting in which micron-scale structures need to be characterized comprehensively in 3 dimensions over mm to cm scale.

## 1. Introduction

### 1.1 Challenges in Mesoscale Imaging and Micro-CT

Micro-Computed Tomography (micro-CT) is rapidly becoming a valuable imaging technique for applications where the generation of high-resolution, isotropic, 3-dimensional (3D) datasets is essential for both qualitative and quantitative phenotyping as well as visualization (Weinhardt *et al*., 2018; Hur *et al*., 2018; Babaei *et al*., 2016; Seo *et al*., 2015). Serial electron microscopy offers nm-scale resolution at the cost of field-of-view and data handling, as acquiring and analyzing mm-scale samples is a protracted process (Hildebrand *et al*., 2017). Lower-resolution imaging methods like magnetic resonance imaging generate proportionately smaller datasets for the same field-of-view, suitable for scanning much larger samples, but are limited to tens to hundreds of microns in resolution (Dodd *et al*., 1999; Edlow *et al*., 2019). Micro-CT bridges the gap between resolution, field-of-view, and analytical feasibility by offering isotropic micron-scale resolution for millimeter to centimeter-scale samples (Mizutani *et al*., 2013; Ding *et al*., 2019). Such images have become a prioritized need in the disease model research community, to enable the anchoring of spatially resolved large scale molecular analyses (Weinhardt *et al*., 2018). Large-field images of sufficient resolution may potentially enable the quantitative measurement of pathological tissue features that differentiate between pathological processes that are divergent in terms of clinical outcome, but indistinguishable in histological (2D) analysis. Commercial micro-CT scanners available from companies such as Zeiss, Siemens, and Bruker use either commercial flat-panel detectors or microscope objective lenses with field-of-view–to–resolution ratios of about 1,000, as opposed to the 10-fold increase demonstrated by our work.

### 1.2 Advances in Synchrotron-Based Micro-CT

Many efforts to improve the resolution and field-of-view of micro-CT have focused on larger detector arrays and non-traditional image acquisition/reconstruction techniques for use in synchrotron facilities. Synchrotron sources’ tunable, brilliant, monochromatic, and parallel X-rays also provide an opportunity for the development of improved optics systems such as the one demonstrated here. Image acquisition and data processing techniques such as phase-contrast imaging for unstained soft tissue, spiral-CT acquisition for increasing field-of-view, dual-energy scanning for simultaneous use of multiple stains, and post-acquisition software-based processing tools such as distortion correction have allowed micro-CT imaging to mitigate the limitations in detector technologies (Barbone *et al*., 2021; Martins de Souza E Silva *et al*., 2017; Sawall *et al*., 2012; Vo *et al*., 2015; Pelt and Parkinson, 2018). Recently, a phase-contrast application using a 2,000 × 2,000 PCO edge 4.2 CLHS camera was able to achieve cm-scale whole-organ imaging at 25 μm voxel size with the capability of zooming in to 1.4 μm voxel size in areas of interest. This was achieved through a combination of benefiting from the increased coherence and brilliance of a 4G synchrotron, a custom sample mounting and image acquisition approach to extend the dynamic range of their two detectors, and stitching/feathering for reconstructions (Walsh *et al*., 2021), Vescovi *et al*., 2018). Alternatively, the development of higher-resolution optics systems allows for their application in any of the previously mentioned applications; one research group adapted a commercial 36-megapixel digital single-lens reflex camera for visualizing whole secondary pulmonary lobules in large human lung specimens, resulting in a 40.6 mm wide × 15.1 mm high field-of-view with 3.07 μm pixel size without any advanced acquisition techniques (Umetani *et al*., 2020). Despite these advances in detectors and acquisition/reconstruction mechanisms, there is still a fundamental trade-off between resolution and field-of-view in micro-CT systems; cm-scale field-of-view applications are generally limited to a multiple micron resolution, while the detection of micron-scale features is normally limited to sub-mm fields-of-view (Bailey *et al*., 2017; Kc *et al*., 2019; Shearing *et al*., 2011). This poses a challenge to studies seeking to thoroughly investigate large, complex samples at high resolution, ranging from disease models (e.g. developing mice, zebrafish, *Daphnia*), to micro-circuits in electronic components, to fuel spray systems (Wong *et al*., 2012;Ding *et al*., 2019; De Samber *et al*., 2008; Lall and Wei, 2015; Tekawade *et al*., 2019).

### 1.3 Development of the Wide-Field Detector

Most detector systems for micro-CT applications use objective lenses designed for cameras with sensors with up to 5 million pixels. These systems have been limited by both the availability of scientific CMOS sensors and field of view limitations of magnifying optics. The rapid development of CMOS sensors for both consumer and commercial imaging applications has led to the availability of CMOS sensors with over 100 million pixels with excellent noise performance at reasonable cost levels. The key to making use of these large-format sensors in a detector system with micron-scale resolution is an objective lens with large field-of-view, low magnification, high numerical aperture, low distortion, and high-throughput for the scintillator emission spectrum. Since no commercial lens currently meets these requirements, we collaborated with Mobile Imaging Innovations, Inc. to develop a custom objective lens that will match an Advanced Photo System (APS)-size CMOS sensor. In our prototype we used a 71 megapixel thermo-electrically cooled CMOS camera in 10,000 × 7,096-pixel format and 3.1 μm pixel size. The custom objective lens provides 6.2x magnification and switchable numerical aperture of 0.6/0.75, with diffraction-limited resolution achieved across the entire field at 0.6. Effectively this objective lens provides the field of view of a conventional 4x microscope objective lens and the resolution of a 40x objective. Our custom detector represents a factor of 100x gain in field-of-view and 1,000x gain in scan volume compared with commercial objective lenses, enabling far more extensive studies of complex material science samples, engineered devices, and biological systems of mm to cm scale. Here, we present measurements of our detector’s resolution and field-of-view based on data acquired at the Berkeley Advanced Light Source (ALS) synchrotron with a test phantom consisting of parallel lines and a Siemens star with micrometer line width, and two biological species –*Daphnia magna* and *Danio rerio* (zebrafish). This system sets a new benchmark for what is attainable in micro-CT detector technologies and can be implemented in both synchrotron and tabletop-based micro-CT facilities.

## 2. Materials and methods

### 2.1 Detector Design

All experiments conducted at beamline 8.3.2 reported in this manuscript used LuAG:Ce scintillators with a 15 mm diameter and 100 μm thickness from Epic Crystal, polished on both sides with anti-reflective coating applied on the exit side. Our custom objective lens was developed by Mobile Imaging Innovations, Inc, and CMOS cameras were acquired from Vision Systems Technology. The camera used in this experiment is an APS-sized CMOS sensor (CHR70M) with 71 million pixels arranged in 10,000 × 7,096-pixel format with 3.1 μm pixel size, resulting in a 5 mm × 3.5 mm field-of-view (Vieworks VP-71MC produced by Vision Systems Technology). Although this is not a designated scientific CMOS sensor, we used a Grade-A chip with 3-stage thermal electric cooling to significantly improve the noise performance and reduce the number of bad pixels. The resulting performance approaches that of a scientific-grade CMOS sensor.

This custom lens provides 6.2x magnification with a switchable numerical aperture of 0.6/0.75. The diffraction-limited optical resolution of 0.5 microns is achieved at an NA of 0.6 across the entire field-of-view and maintained when used at 0.75 with a 55% increase in throughput. It achieves an image circle of over 6 mm on the scintillator plane and an effective field-of-view of 5 mm x 3.5 mm and pixel size of 0.5 microns with this CMOS sensor. The distortion is under 2%. To our knowledge, the only optics with comparable resolution and field-of-view are projection lenses for lithography, which operate at a much narrower bandwidth at much greater cost (Kawata *et al*.1989).

### 2.2 Animal Husbandry and Sample Preparation

*Daphnia magna* (*D. magna*) were raised in modified artificial freshwater “Aachener Daphnien Medium” or ADaM (Ebert, 1998) at room temperature (20°C ± 1°C) under a 16-hour light/8-hour dark photoperiod. The animals were fed three times weekly with 3.0 × 107 cells/mL of green microalgae (*Raphidocelis subcapitata*, previously known as *Selenastrum capricornutum*) and 0.1 mg/mL of dissolved bakers’ yeast. The embedding protocol was adapted from Lin *et al*., (2018). *D. magna* were fixed in Bouin’s for 2 days at room temperature and then stained with 2% PTA (in 70% ethanol) for another 2 days prior to embedding.

Zebrafish were housed in Penn State Zebrafish Functional Genomics Core recirculating aquaria at an average temperature of 28°C and on a 14-hour light/10-hour dark cycle. The fish were fed three times a day with a diet consisting of brine shrimp and dry flake food. The wildtype zebrafish were staged according to a standard developmental staging series (Kimmel *et al*., 1995). The zebrafish were subjected to the same fixation protocol which was described by Lin *et al.*. (2018). All procedures on zebrafish were approved by the Institutional Animal Care and Use Committee (IACUC) at the Pennsylvania State University College of Medicine, ID: PROTO201800300, “Developing Tissue-, Cell-, and Protein-specific Staining for histo-tomography, a form of X-ray Microtomography (micro-CT)” and PRAMS201445659, “[Groundwork for a Synchrotron MicroCT Imaging Resource for Biology (SMIRB)](https://iacuc.psu.edu/IACUC/sd/Rooms/DisplayPages/LayoutInitial?Container=com.webridge.entity.Entity%5BOID%5B3CC716D490D97E43AFA79C000422842D%5D%5D).”

### 2.3 Imaging Parameters

All samples were scanned at Beamline 8.3.2 at the Advanced Light Source at Lawrence Berkeley National Laboratory. Alignment of the detector with the synchrotron beam was accomplished with a custom mounting system comprised of two horizontal Newport MTN100PP stages, a vertical Newport IMS150PP stage, Thorlabs 90-degree mounts, and custom-designed milled adapters to integrate imperial and SI components. Scan geometry was recorded as 7.6 cm between the object and scintillator after orthogonal alignment to the beam for QRM Nanobar phantom (QRM) acquisitions, and 4.3 cm for *Daphnia*. All scans of the phantom and *Daphnia* were acquired at 20 keV, as a sequence of 150 ms projections for *Daphnia* and 5000 ms for the phantom, due to the phantom scans being acquired at lower flux during the Beamline’s operation on 2-bunch mode. The QRM Nanobar Phantom was acquired as 7873 projection images. Projections were acquired over 180 degrees and reconstructed with parallel-beam reconstruction techniques through TomoPy (Fig. 3). The number of projections was determined by

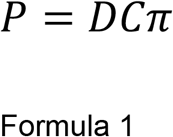

where P is the total number of projections, D is the furthest horizontal distance in pixels from the center of rotation to the edge of the sample in the field-of-view, and C is the number of pixels per μm. *Daphnia* scans were acquired with ~3000 projections each, depending on the diameter of the sample.

### 2.4 Detector Setup at Beamline 8.3.2

We performed the initial detector test and validation at the Berkeley ALS 8.3.2 beamline; synchrotron beam properties offer advantages for testing detector systems compared to commercial tabletop sources (Fig. S1): the brilliant X-rays allow for shorter scan times and as a result reduced drift artifact in images (Fig. S1A, B). The high monochromaticity, tunability, and coherence aspects of the beam allow us to measure the detector’s response as a function of X-ray energy and selectively tune phase contrast edge enhancement, as well as simplify the reconstruction process with conventional parallel-beam reconstruction approaches (Fig S1C, D). Single CT dataset acquisitions took approximately 30 minutes at 20 keV and an object-to-scintillator distance of 7.6 cm, compared to >48-hour imaging sessions under similar conditions to achieve a comparable signal-to-noise ratio (SNR) on a local source. In order to efficiently utilize our detector at the ALS 8.3.2 beamline as well as for easy adaptation to either other synchrotron facilities or local desktop applications, a motion control system provides 3 axes of motion with a 10 cm × 10 cm × 15 cm range to align the detector to the beam (Fig. 1).

**Figure 1:**
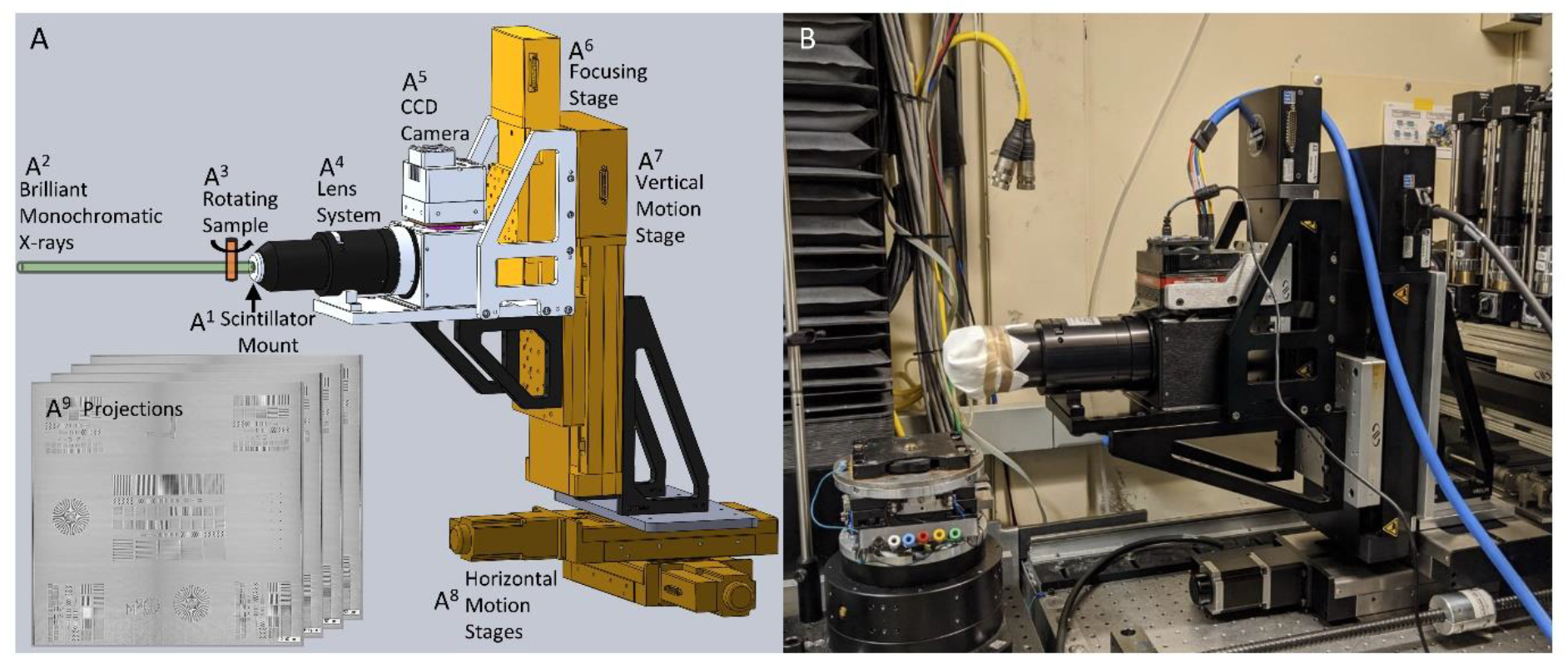
Generalized beam alignment and image acquisition setup for application in synchrotron facilities and tabletop sources. Our custom detector operates using a scintillator (A^1^) to convert x-rays (A^2^) passing through a sample (A^3^) into visible light. The fluorescence image is then focused by the objective lens (A^4^) onto the CMOS sensor surface (A^5^). Commercially available Newport stages (A^6^-A^8^) are utilized to adjust the camera relative to the focal point, as well as to translate the entire detector through 3D space (15 cm vertical travel and 10 cm horizontal travel in any direction) for optimal alignment with any synchrotron or tabletop generated beam. Projections (A^9^) acquired by the detector are saved using the Harvester Python library and later reconstructed with the TomoPy toolbox in Python. The actual setup as it was used in the Berkeley ALS 8.3.2 beamline hutch is shown (B) in comparison to the CAD design.

### 2.5 Acquisition, Reconstruction and Data Visualization

Image acquisition from the CMOS camera was controlled by a Python script using the Harvester library (https://github.com/genicam/harvesters). The stage holding the sample was set to rotate at constant velocity and trigger camera-based projection acquisition at preselected intervals. Flat field correction, ring artifact reduction, and reconstruction were performed using the TomoPy toolkit (Gürsoy *et al*., 2014). Flat field correction used either the flat image taken before or after sample scanning, depending on which time point yielded the most contrast with test projections. Reconstructions were computed using the gridrec algorithm (Dowd *et al*., 1999; Rivers, 2012) with a Shepp-Logan filter. Stripe correction on the sinogram domain was performed using a combination of wavelet–Fourier filtering and Titarenko’s algorithm (Miqueles *et al*., 2014; Münch *et al*., 2009). Reconstructions resulted in a nominal isotropic voxel size of 0.52 μm, as estimated by the number of pixels spanning the 1.03 mm diameter of a reconstructed polyimide tube used for sample embedding. All data was visualized in ImageJ.

### 2.6 Resolution Measurements

All visual confirmation of resolution based on demarcations present on the phantom, whether in projection or reconstruction domains, was done by visualizing data in ImageJ and adjusting image window/level to detect contrast. Intensity line profiles were calculated as a vector of individual intensity values taken across indicated areas. The line spread function (LSF) was calculated as

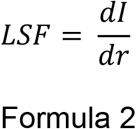

where I is the intensity line profile and r is distance. The modulation transfer function (MTF) was then calculated as the discrete normalized Fourier transform of L in the direction of intensity variation of I.

## 3. Results

### 3.1 Characterization of Field-of-View

Upon centering the beam with the detector, initial projection tests confirmed our detector’s field-of-view as larger than what is available for typical synchrotron applications (Fig. 2A). While the detector’s 5 mm horizontal field-of-view was sufficiently covered by the beam, the 3.5 mm vertical range encapsulated the full beam height, allowing the acquisition of samples previously unattainable on similar setups (Fig. 2B). The beam height at 8.3.2 for these experiments was limited by the monochromator—using longer multilayer mirrors in a redesigned monochromator would allow us to fully utilize the field of view of this detector. The 8.3.2 beamline’s optics capabilities can expand the horizontal field-of-view of their detection system to >10 mm at a 9 μm pixel size. For comparison, the beamline’s 2000 × 2000-pixel CMOS camera set to a 0.5 μm pixel size would have a field-of-view of 1mm × 1 mm, 1/17.5 of our detector’s area (Fig. 2A, B, C). As a functional demonstration of the field-of-view, we imaged a juvenile, 57-day old zebrafish (Fig. 2B). A 2000 × 2000-pixel detector with an equivalent 0.5 μm pixel size would require hundreds of individually acquired and stitched scans across both the horizontal and vertical field-of-view to cover the same reconstructed volume as 4 scans on the custom detector stitched only in the axial dimension (Fig 2C). When performing tomography of large objects, stitching data in the axial direction is relatively straightforward and can be done after tomographic reconstruction. Stitching data in the radial direction is challenging since this must be done in the sinogram domain, and small inconsistencies in the stitching or calibration of rotation axes will be amplified by tomographic reconstruction and lead to artifacts in reconstructed images (Vescovi *et al*. 2018).

**Figure 2:**
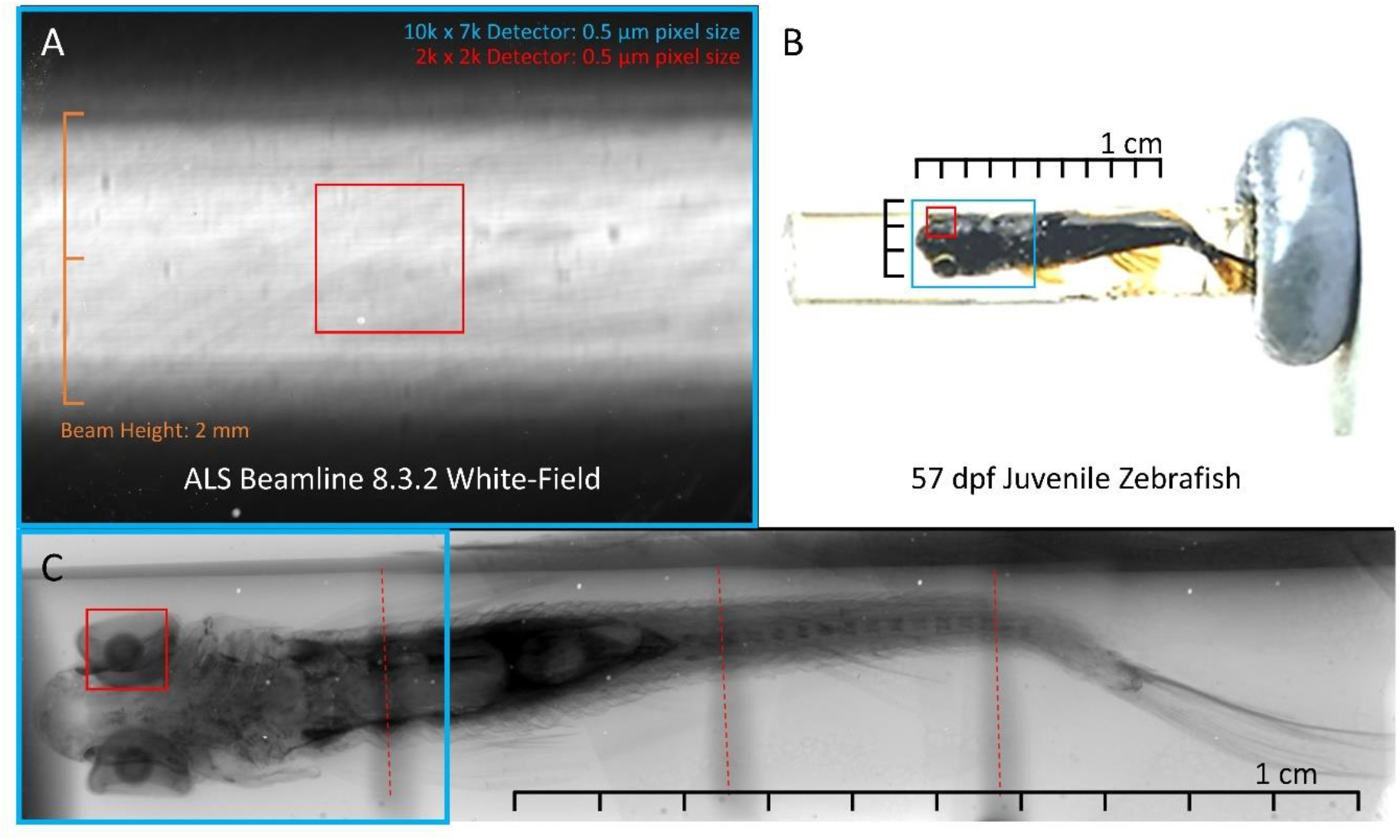
Field-of-view characterization utilizing ALS facilities synchrotron beam and large-scale biological sample. The entirety of the beam at the ALS 8.3.2 beamline is completely encapsulated by the detector’s field-of-view in the vertical direction while maintaining 0.5 μm pixel size (A). A 2000×2000 pixel field-of-view at the same 0.5 μm pixel size is shown in red. To illustrate the field-of-view in a more biologically applicable and practical sense, we present a cm-scale, juvenile, 57 day-post-fertilization (dpf) zebrafish embedded in resin and stained with phosphotungstic acid (PTA) (B), with the same 2,000 x 2,000 pixel array in red and our detector’s field-of-view in blue. Rotating the camera by 90 degrees makes it possible to scan in a single view horizontally and stitch in 4 sections vertically, compared to approximately 500 scans on a 2,000 x 2,000 detector to cover the same volume (C). Projections were acquired using a NOVA 96000 local source to demonstrate field-of-view, and the 2000 x 2000 pixel field-of-view is again shown in red.

### 3.2 Quantification of Resolution

Our resolution was qualified using a Nanobar phantom fabricated by QRM consisting of 2 perpendicular 2.96 mm^2^ silicon dies that each include sets of specified line pairs and dot patterns ranging from 10 μm to 1 μm feature size, Siemens stars converging to 1 μm at the points, and an L edge, encapsulated in a 5 mm thick solid plastic (Fig. 3). All structures are etched in the silicon die with a recorded depth of 5-15 μm. The field-of-view offered by our custom detector enabled the capture of both chips in a single acquisition sequence (Fig. 3A).

**Figure 3:**
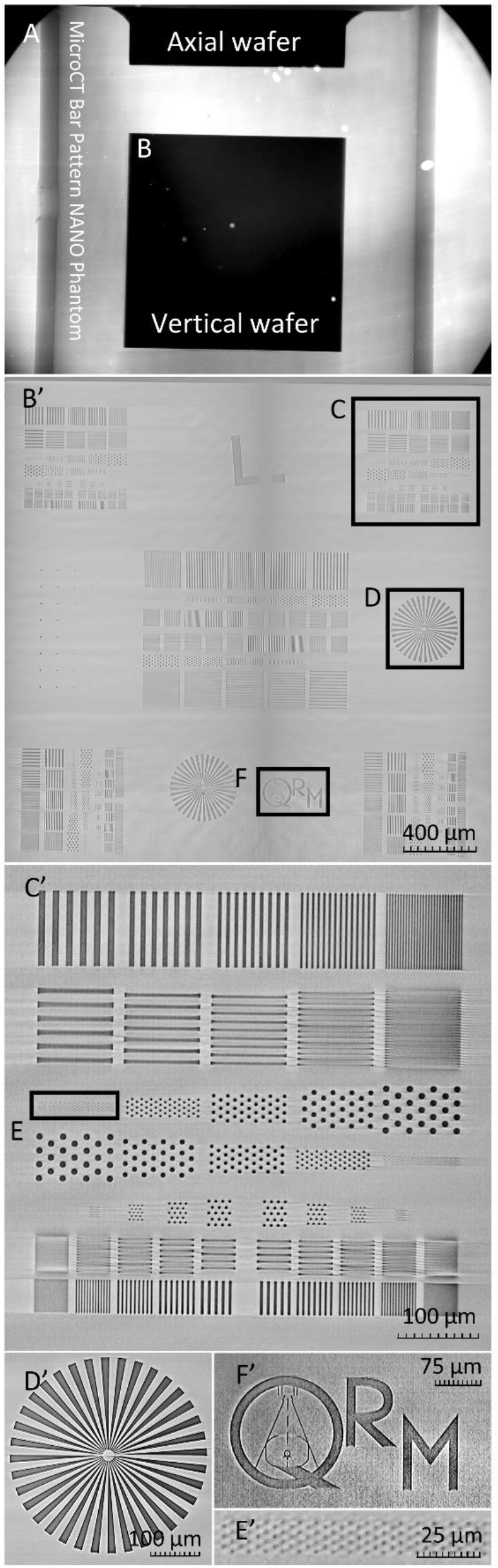
Reconstruction of QRM nanobar phantom die acquired over a single projection series. After showing a projection encapsulating both case die (A), We present multiple zoomed-in elements of the reconstructed vertical die (B’) that qualitatively and unequivocally demonstrate our resolution via elements of defined size. A pattern cluster containing line pairs and dots ranging from 10 μm to 1 μm (C’) is emphasized, along with a dot pattern containing elements that are 2 μm in diameter (E’). The etched QRM symbol (F’) is also shown along with the reconstruction of the same Siemens star (D’) previously shown as a projection in Fig. S1.

The spatial resolution of the 3D reconstruction was measured from a segment of the reconstructed image that contains line pairs as the simplest method to showcase the detector’s ability to discern 1 μm features (Fig. 4). Although the detector provides a nominal resolution of 0.5 μm, the width of the smallest 1 μm line pairs is at the Nyquist-limited resolution with our presented acquisition and reconstruction pixel size of 0.5 μm (Fig. 4).

**Figure 4:**
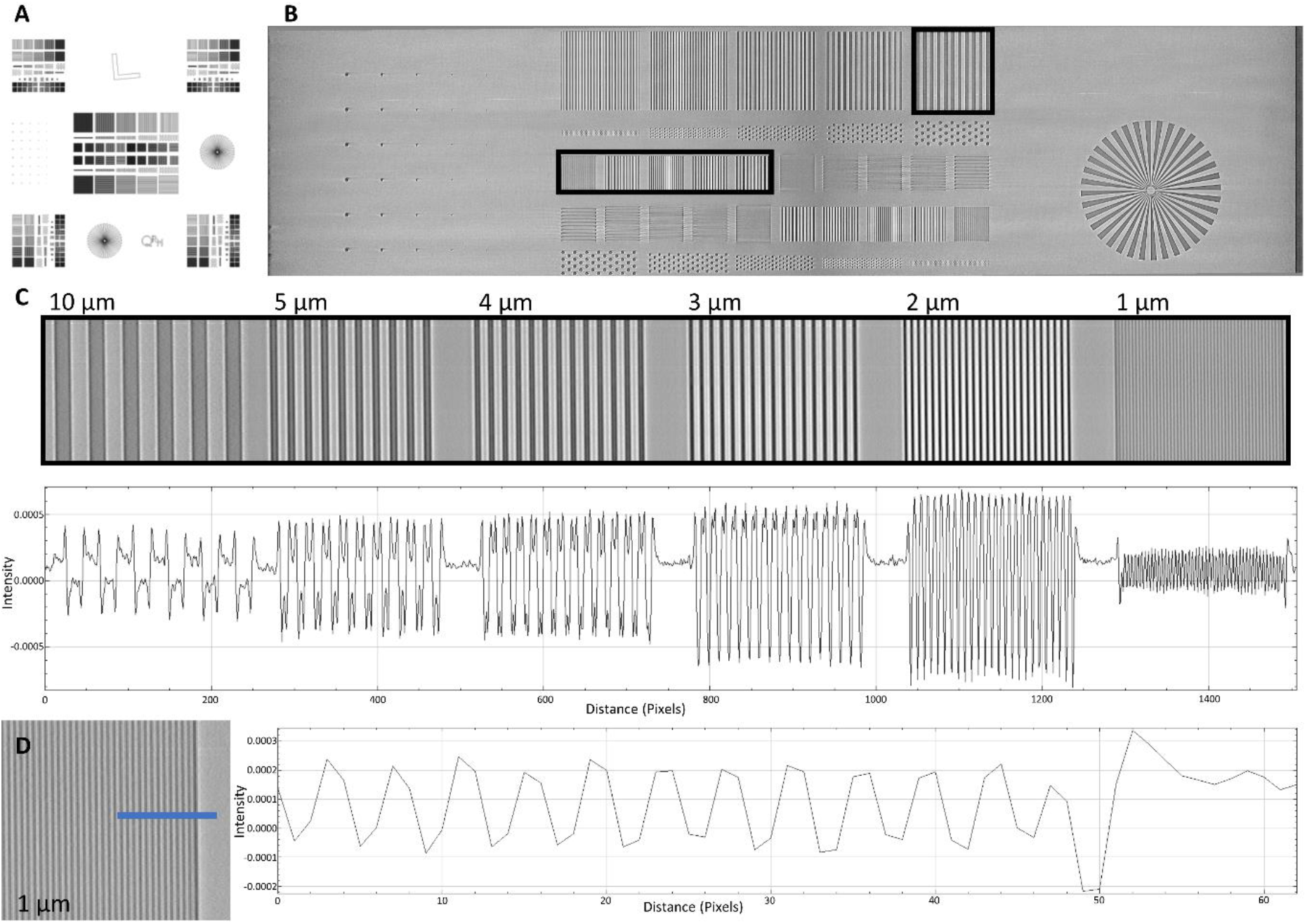
Verification of resolution utilizing a QRM bar pattern phantom. A QRM-manufactured phantom including line and point patterns, siemens stars, and an L edge (A) was scanned at the Berkeley ALS and locally reconstructed to characterize the resolution of the combined lens system and camera detector. A portion of the reconstructed phantom is presented (B) showcasing line pairs ranging from 10 to 1 μm in width. A line profile of the indicated regions is shown, validating resolution up to 1 μm (C). A smaller line profile taken across the 1 μm line pairs that highlights signal shape is shown (D). Signal at the edges of line pairs is enhanced by phase effects caused by the silicon-air interface. Pixel counts between line pairs confirm reconstruction at 0.5 μm, as each line of the 1 μm pattern is 2 pixels thick.

An intensity profile taken through these line pairs confirmed visible contrast down to a resolution of 1 μm. As an additional measurement of our resolution, the L edge on the reconstructed scan of the Nanobar phantom was used to calculate the line spread function (LSF) as well as the modulation transfer function (MTF) in both the projection as well as the reconstruction domain. 400 projections acquired at a single angle were averaged for this measurement, with the full width at half maximum of the LSF found to be 1.183 μm, and a drop-off to the 0.1 amplitude of the MTF at 0.85 line pairs per μm (Fig. 5).

**Figure 5:**
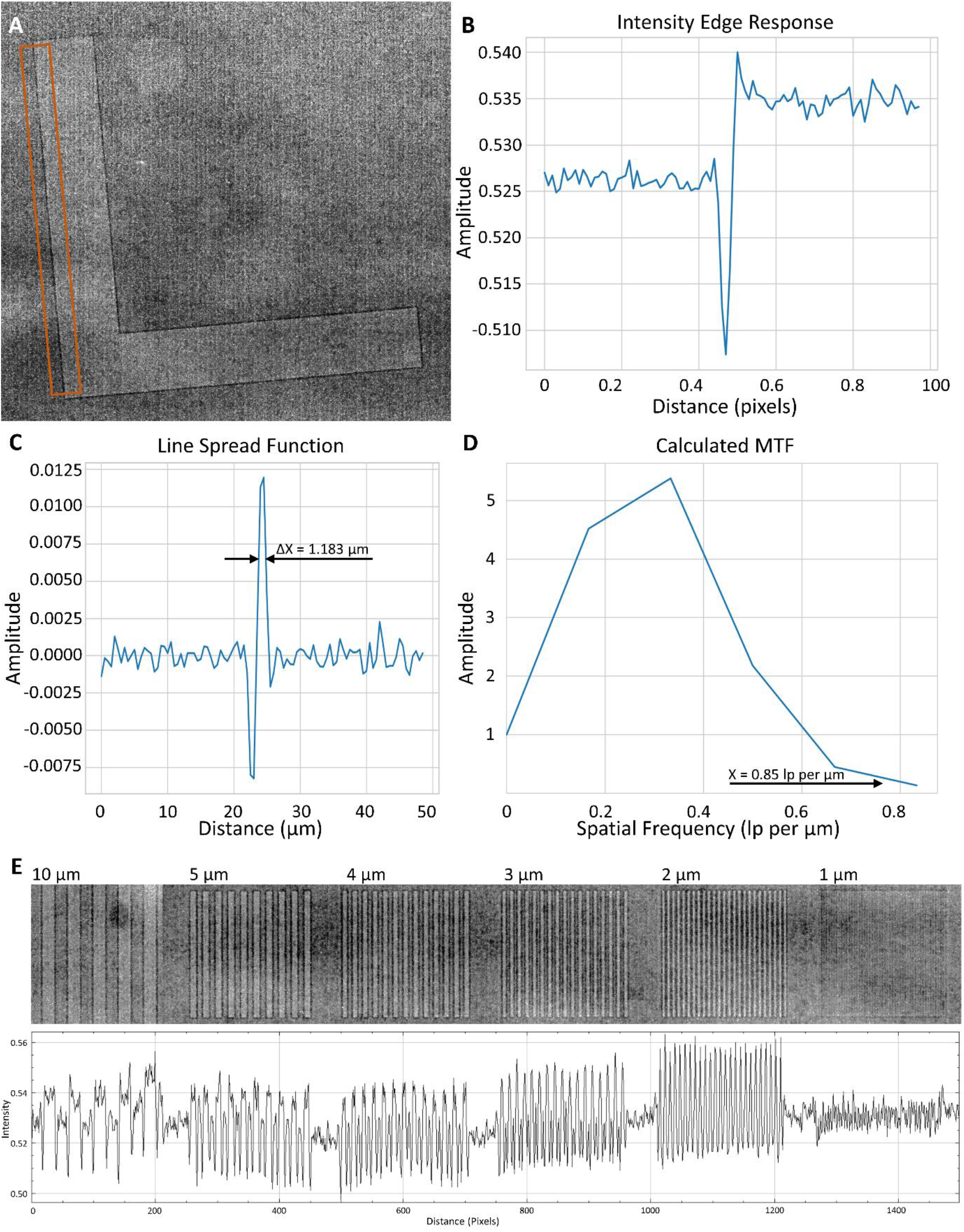
Slanted Edge Response and calculation of Modulation Transfer Function (MTF) in the projection domain. A 400 projection series taken at the same angle of the indicated area of the slanted L edge (A) was used to record the associated intensity line profile (B). The Line Spread Function (LSF) (C) was taken as the first derivative of B across pixel distance, and the MTF (D) was calculated as the discrete normalized Fourier transform of the portion of C covering the edge. The full width at half maximum amplitude of the LSF was recorded at 1.183 μm (C), with the indicated 0.1 MTF amplitude located at 0.85 line pairs per μm (D). Line pairs are presented with a corresponding intensity line profile indicating contrast within the 10, 5, 4, 3, 2, and 1 μm line pair groups (E). Edge enhancement can be visualized easiest in the intensity profile region corresponding to the 10 μm demarcations.

These results are consistent with our ability to qualitatively discern 1 μm features on the phantom at a 0.5 μm pixel size. Overall, we achieve a pixel size of 0.5 microns and a spatial resolution of ~1.2 microns as measured at an NA of 0.6 across a 5 mm × 3.5 mm field-of-view, and these specifications are maintained when used at NA of 0.75 with a 55% increase in throughput. The effects of tomographic reconstruction, although boosting the signal through slices of the phantom used for analysis, increased the full width at half maximum amplitude to 1.575 μm and the 0.1 MTF amplitude to 0.278 line pairs per μm (Fig. S2). This slight loss in resolution in the reconstruction domain is likely due to the various interpolations and filters used during tomographic reconstruction and post-reconstruction interpolation due to rotating in each of three orthogonal planes to bring the test pattern *en face*.

### 3.3 Biological Applications in *Daphnia*

*Daphnia magna* is a freshwater crustacean that plays a significant role in the food web, has relatively high sensitivity to environmental contaminants, and has been commonly used as a model organism for toxicity testing and environmental biomonitoring for over half a century (Anderson, 1944; EPA, 2002, 2016; OECD 2001, 2004, 2012). To demonstrate the field-of-view and resolution of our detector system, we scanned an entire adult wild-type *D. magna* at the ALS. The resulting 3D reconstructions allow virtual, histology-like “sections” in any plane and 3D rendering at user-defined scales to reveal microanatomic features of organ systems, tissues, and cells in context of the entire organism at sub-cellular resolution (Fig. 6). Details of thicker tissues or organs, such as the connections between ommatidia, optic nerves, optic lobe and brain (Fig. 6C), and the long setae (hair-like structures) of filter plates (Fig. 6D) can be visualized through customizing maximum intensity projection (MIP) thicknesses. Reconstructed voxel sizes of 0.5 μm allow visualization of cellular structures, such as nucleoli (about 2 μm in diameter) of gut epithelial cells (Fig. 6E), nucleoli of fat cells (Fig. 6F), and yolk globules in a developing embryo (Fig. 6G). Visualizing and characterizing changes in these structures are important because they play important roles in development, reproduction, and energy processing (Fryer, 1991, Quaglia *et al*., 1976, Zaffagnini and Zeni, 1986; Colbourne *et al*., 2011).

**Figure 6:**
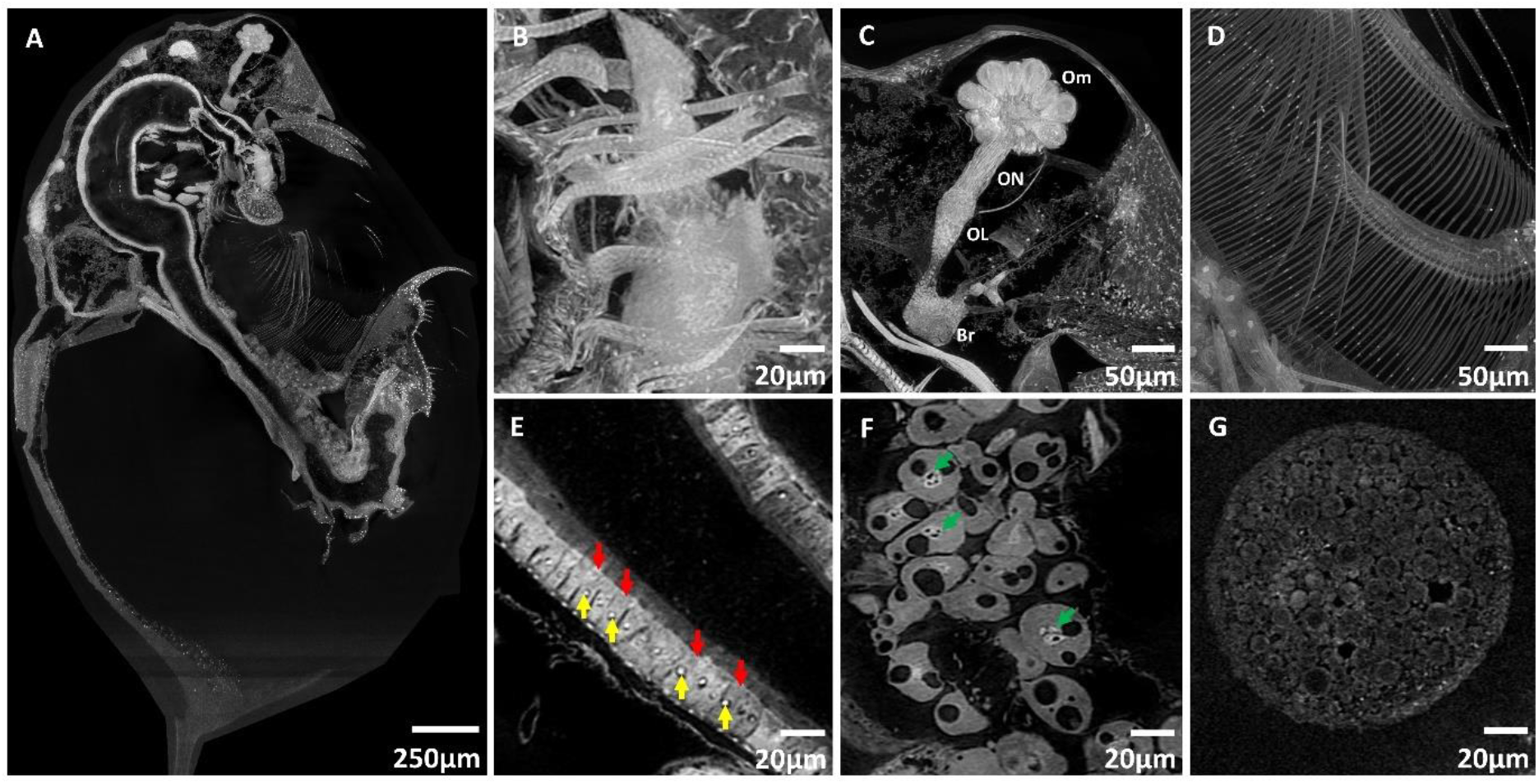
Adult D. magna stained with PTA showing sub-cellular resolution for different cell types and structures. **(A).** Cell types and structures that can be visualized by customizing various stack thicknesses included: muscle striations (about 2 μm) on the labral muscles (B), ommatidia (Om), optic nerves (ON), optic lobe (OL) and brain (Br) (C), setae of the filter plates (D), nucleoli (yellow arrows - ~ 2 μm diameter) in the gut epithelial cells (red arrows) (E), nucleoli (green arrows) in the fat cells (F) and yolk globules (G) in a developing embryo. Panels A to C were generated from maximum intensity projections (MIP) of 50 slices, panel D from a MIP of 100 slices to visualize thicker 3D structures and panel E to G from a MIP of 10 slices. A MIP of 10 slices is equivalent to a 5 μm thick histology slide.

The ability to interrogate the cellular and organ-based structures throughout entire organisms is essential for characterization of toxicity response, development, and morphological effects. To exemplify the detection of cellular change, we imaged an abnormal wild-type *D. magna* where the compound eye had developed outside the carapace (Fig. S3). Micro-CT imaging detected the deformities of the ommatidia (less than the usual 22 ommatidia), longer optic nerves, and absence of an optic lobe (Fig. S3B). Moreover, we detected cellular changes (vacuolation and sloughing of epithelial cells) in the gut (Fig. S3C) which could potentially go unnoticed if using an imaging technology that does not consider the entire specimen at cellular detail, such as histology, electron microscopy, or micro-CT that compromises on either field-of-view or resolution. With our detector, we envision that whole-organism imaging will become a comprehensive approach to systematic characterization of 3-dimensional morphological and cellular changes, not only in *Daphnia* research, but for other mesoscale model organisms.

## 4. Discussion

### 4.1 Detector Capabilities and Nonconventional Applications

The compromise between resolution and field-of-view is a challenge for imaging that has not been fully addressed for mesoscale imaging applications by either synchrotron facilities or commercial companies. Current synchrotron-based high-resolution micro-CT systems rely on microscope objective lenses to magnify scintillated images or flatpanel detectors developed for radiology. Therefore, the resolution to field-of-view is limited to microscopy standards, about 1/1,000. In most sub-micron resolution applications, the field-of-view is typically limited to 1 mm. Our custom detector provides a 1 micron-scale resolution at a 5 mm field-of-view, enabling unique imaging applications of large samples in single field-of-view acquisitions. Additionally, even larger samples falling outside the capabilities of the detector can be scanned by further expanding the wide field-of-view with spiral micro-CT acquisition (Pelt and Parkinson 2018). Furthermore, mosaic spiral-CT, which takes multiple spiral-CT passes, can be implemented to significantly expand the effective field-of-view beyond that of the detector without having to spend resources on stitching a large number of individual small scans in both horizontal and vertical directions, which often propagates stitching errors that create artifacts in the reconstructed 3D image. An object about twice as wide as the camera’s field-of-view horizontally and many times larger vertically can first be positioned to cover the left half, and a spiral CT sequence can be acquired throughout its full length. Then another spiral CT data set is acquired over the right half. The 2D projections from the two data sets acquired at the same angle and vertical positions can then be stitched pairwise to form a larger 2D projection that spans the entire width and length. A mosaic spiral acquisition approach utilized on this detector will enable a new range of applications for imaging larger biological samples such as intact mouse embryos and excised brains, adult zebrafish, and other use-cases for micro-CT centered on geology, electronics, and clinical research. Alternatively, the field-of-view and resolution combination afforded by the detector can be leveraged to increase throughput on high-resolution imaging of smaller specimens by stacking multiple samples in a single acquisition and reconstruction.

### 4.2 Implications for Model System Research

Properties such as cellular and organellar shape and volume, anatomical locations, intercellular relationships, physiology, environment, and state of health are reflected in morphological changes in the micron scale whose meaning is apparent in the context of the whole organism. Using our detector system, we demonstrate visualization and characterization of non-targeted cellular changes enabled by whole-organism imaging of *D. magna*, in a single acquisition. We envision systematic characterization of 3-dimensional morphological and cellular changes to become a more comprehensive approach in evaluation of environmentally and genetically induced morphological changes. Looking forward, comprehensive phenotyping of larger disease and toxicological model organisms, such as adult zebrafish, mouse embryos, and *Xenopus* for quantitative measurements of tissue architecture and cellular features, will contribute to the systematic understanding of pathological changes.

## 5. Conclusions and Future Work

To accommodate larger samples, a second detector designed for a larger field of view is currently being designed to provide a 12 mm image circle. This new version will use a medium-format sensor with 151 million pixels in 14,192 × 10,640 format to provide a field of view of 10 mm × 7.5 mm. The new objective lens provides the same switchable 0.6/0.75 numerical aperture, and diffraction-limited optical resolution of 0.5 μm at NA of 0.6 and maintains the same resolution at NA of 0.75. The effective pixel size will be 0.7 μm at the scintillator plane. Because the optical resolution scale is less than the effective pixel size, a 3-step super-resolution acquisition process with 0.25 μm step size is needed to fully realize the optical resolution. The combination of these two detector systems will provide additional versatility for different sample size vs resolution combinations.

Along with our testing of new technologies in scintillator polishing, beam monitoring, and quantum efficiency improvements in high resolution camera sensors, we anticipate significant improvements to synchrotron facilities utilizing our detector. In an ongoing collaboration with beamline 8.3.2 at the Berkeley ALS, we are working towards a permanent installation of our detector to make this high field-of-view, high resolution imaging available to the full micro-CT user base. The larger field of view greatly increases the breadth of specimen types that can be imaged, including both biological and non-biological samples. The adaptability of such large-field, high-resolution optical systems to other synchrotron and non-synchrotron applications promises a broad potential range of benefits in the biomedical and physical sciences across the global microCT community.

## Supporting information

Supplemental Figures with Legends

## Acknowledgements

The authors are very grateful to Dr. Damian B. Van Rossum for writing the General User Proposal for the LBL Advanced Light Source and for his continued support and intellectual contribution.

