## Supplemental Figures with Legends for "A Wide-Field Micro-Computed Tomography Detector: Micron Resolution at Half-centimeter Scale"

### Supporting information

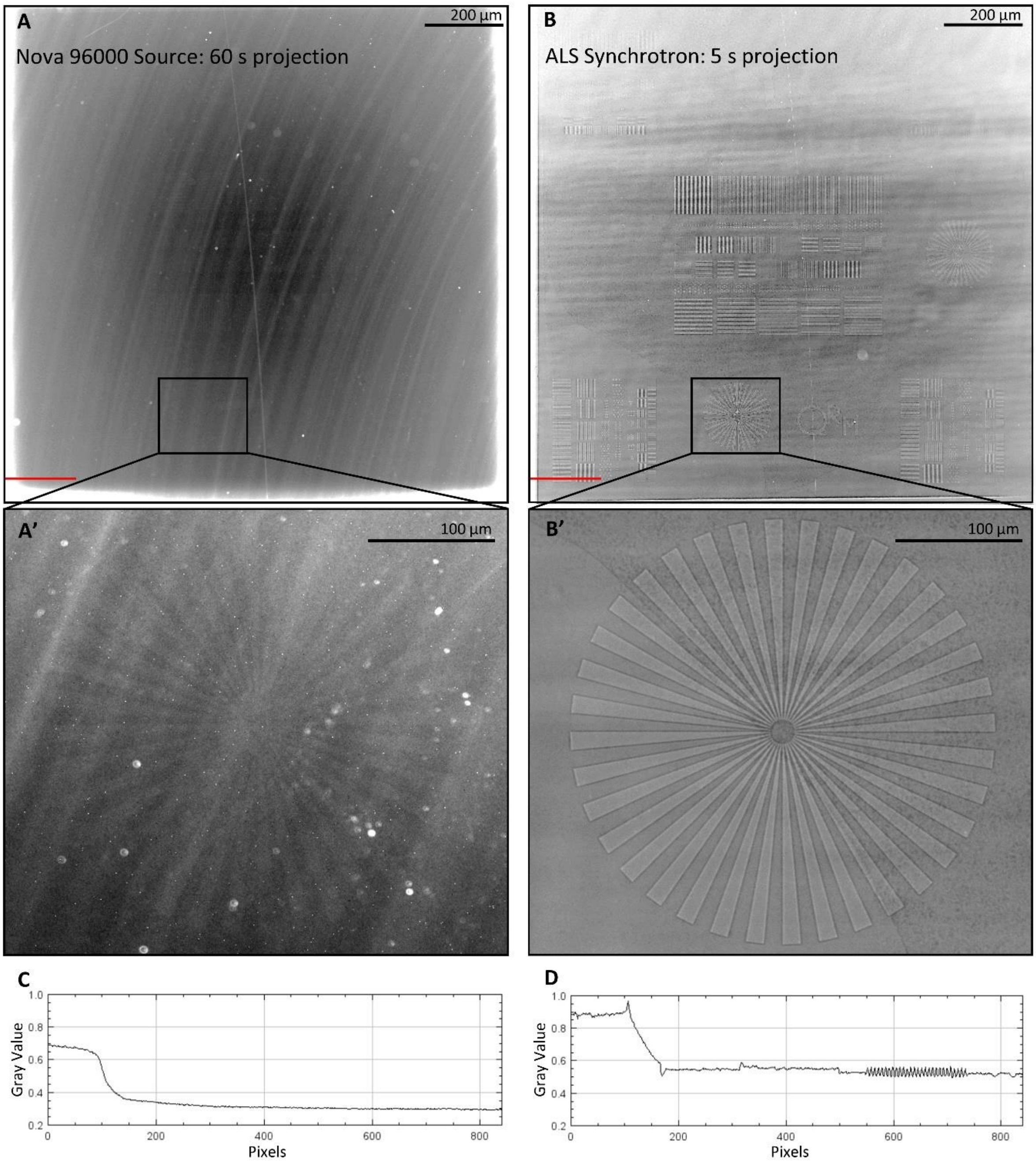


**Figure S1: Projection comparison between the synchrotron beam and a commercially available x-ray source.** Gain-corrected projections of the same QRM nanophantom are presented as acquired from an Oxford Nova 96000 source (A) and the ALS synchrotron (B). Both projections were acquired with the same detector system and scintillator, with the only difference being the beam used for acquisition and exposure time. The monochromatic, parallel synchrotron beam enhances phase effects, allowing the contrast-based observation of features etched into plastic on the phantom, shown in the zoom-ins of A and B as the Siemens star wedges that start at 17 μm and narrow down to 1 μm at the tip (A’, B’). These features are less visible in the same area of the phantom when acquired using our local, cone-beam source (A’). The brilliance of the synchrotron beam also drastically reduces acquisition time per individual projection (150 ms for synchrotron and 60 s for the local source) to achieve comparable SNR, as shown in the line patterns (C,D) indicated by the red lines in A and B.

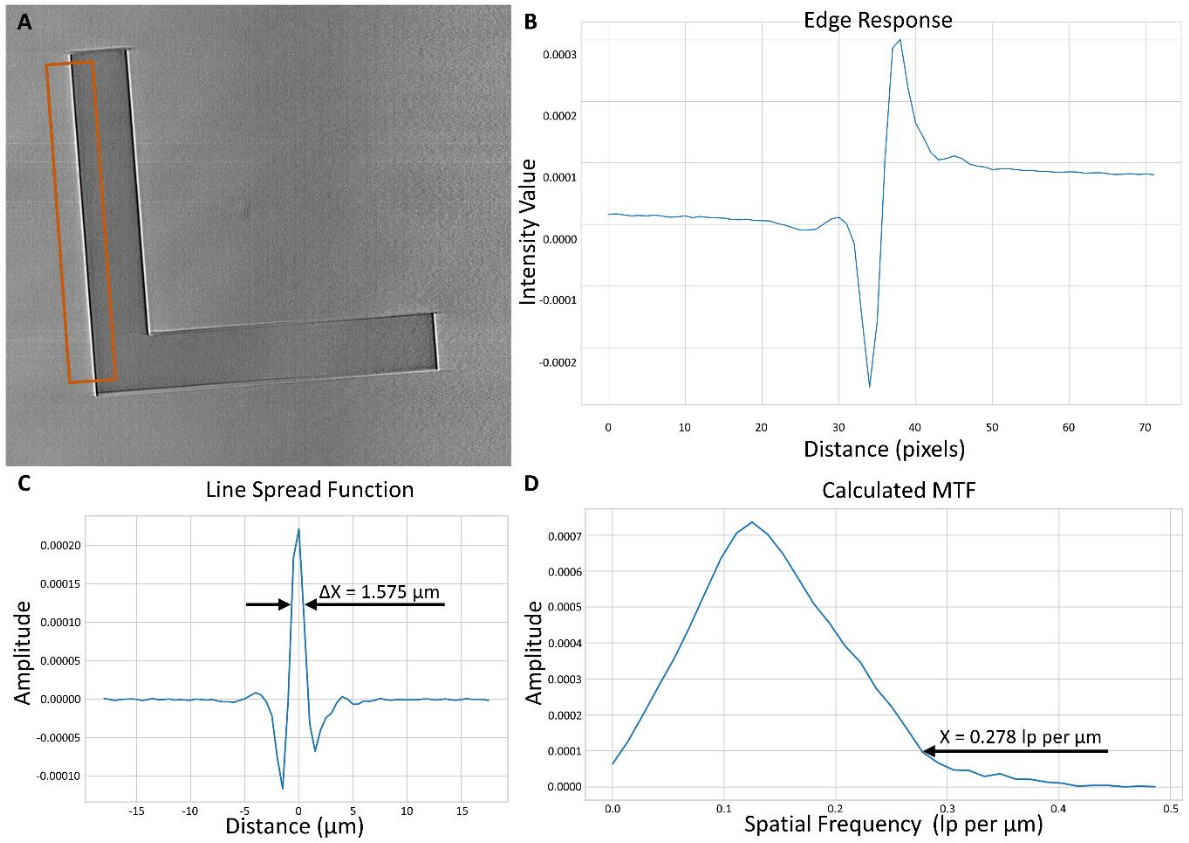


**Figure S2: Slanted Edge Response and calculation of Modulation Transfer Function (MTF) in the reconstruction domain.** The indicated area of the slanted L edge provided by the QRM bar pattern NANO phantom (A) was used to record the associated edge response (B), Line Spread Function (LSF) (C), and MTF (D). The effects of edge enhancement from the Berkeley ALS beam can be observed in the large intensity peaks of the edge response, the minima of the LSF, and decreased MTF at low spatial frequencies. The full width at half maximum amplitude of the LSF was recorded at 1.575 µm, with a 0.1 MTF amplitude at 0.278 lp per µm.

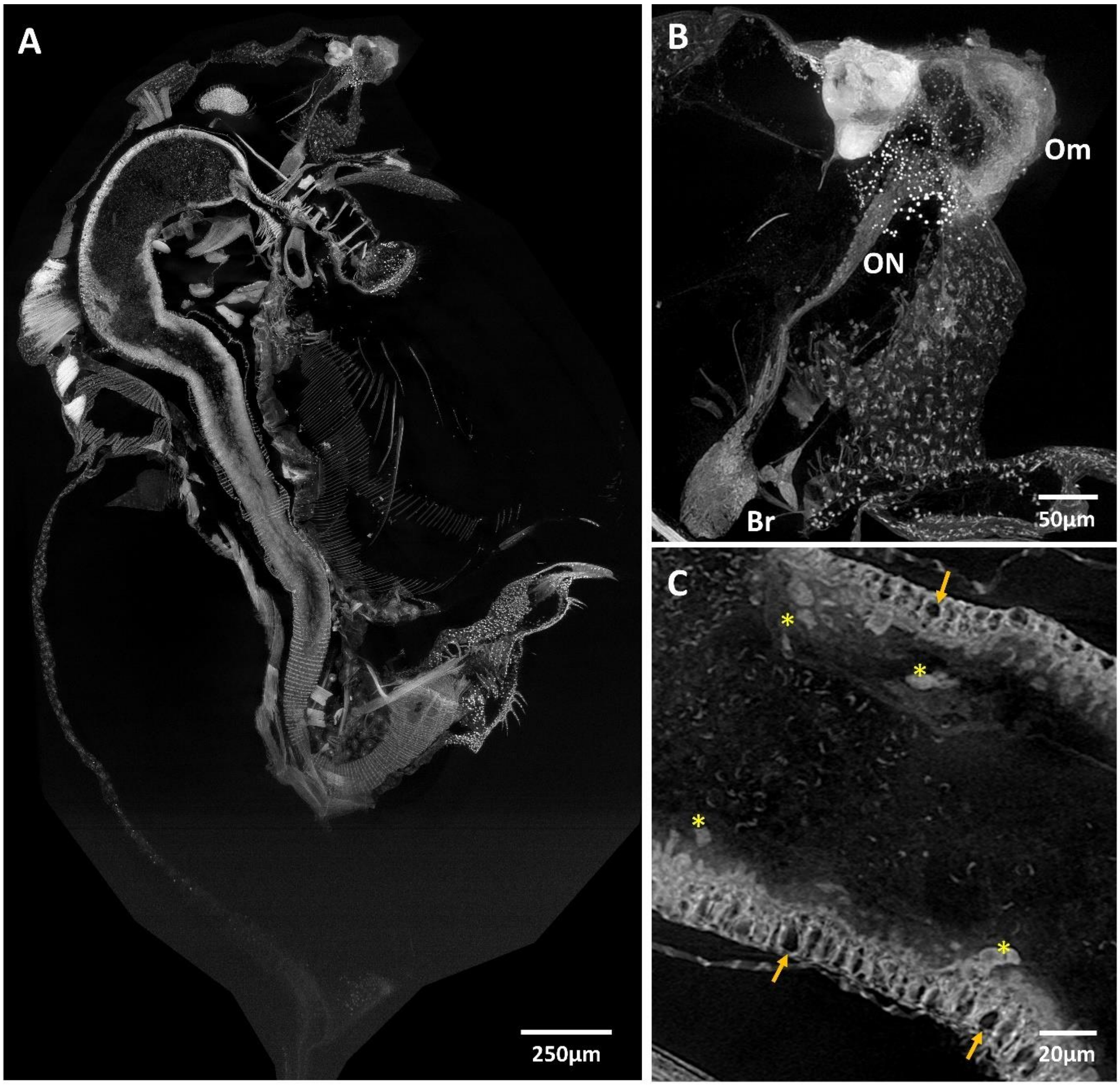


**Figure S3: An unhealthy *Daphnia magna* found in our wildtype population with its ommatidia developed outside the carapace (A).** MicroCT imaging revealed deformities of the ommatidia (Om), longer optic nerves (ON) that connect directly to the brain (Br) but without a corresponding optic lobe (OL) (B). Whole-organism phenotyping also revealed cellular changes in the digestive tract, with vacuolation (orange arrows) and sloughing of gut epithelial cells (asterisks) (C). Panels A and B were generated from maximum intensity projections (MIP) of 50 slices, panel C from a MIP of 10 slices.
